# HPMCP-coated microcapsules containing the Ctx(Ile^21^)-Ha antimicrobial peptide reduces the mortality rate caused by resistant *Salmonella* Enteritidis in poultry

**DOI:** 10.1101/2021.03.29.437537

**Authors:** Cesar Augusto Roque Borda, Larissa Pires Pereira, Elisabete Aparecida Lopes Guastalli, Nilce Maria Soares, Priscilla Ayleen Bustos Mac-Lean, Douglas D’Alessandro Salgado, Andréia Bagliotti Meneguin, Marlus Chorilli, Eduardo Festozo Vicente

**Affiliations:** São Paulo State University (Unesp), School of Agricultural and Veterinarian Sciences, Jaboticabal, São Paulo – Brazil. 14884-900; São Paulo State University (Unesp), School of Sciences and Engineering, Tupã, São Paulo – Brazil. 17602-496; Poultry Health Specialized Laboratory, Biological Institute, Bastos, São Paulo – Brazil. 17690000; São Paulo State University (Unesp), School of Pharmaceutical Sciences, Araraquara, São Paulo – Brazil. 14801-902

**Keywords:** AMP, HPMCP, chicks, microencapsulation, mortality rate

## Abstract

The constant use of synthetic antibiotics as growth promoters can cause great bacterial resistance in chicks. Consequently, the use of these drugs has been restricted in different countries. Antimicrobial peptides have gained relevance in recent years due to their minimal capacity for bacterial resistance and does not generate toxic residues that harm the environment and human health. In this work, Ctx(Ile^21^)-Ha antimicrobial peptide was employed, due to its previously reported largely antimicrobial potential, to evaluate its application effect in laying chicks challenged with *Salmonella* Enteritidis, resistant to nalidixic acid and spectinomycin. For this, Ctx(Ile^21^)-Ha was synthesized, microencapsulated and coated with hypromellose phthalate (HPMCP) to be release in the intestine. Two different doses (3.2 and 6.4 µg of Ctx(Ile^21^)-Ha per kg of isoproteic and isoenergetic poultry feed) were included in the chicks’ food and administered for 28 days. Antimicrobial activity, effect and response as treatment were evaluated. Statistical results were analyzed in detail and indicate that the formulated Ctx(Ile^21^)-Ha peptide had a positive and significant effect in relation to the reduction of chick mortality. There was a significant difference in laying chick weight between the control and microencapsulation treatment groups as a function of time. Therefore, the microencapsulated Ctx(Ile^21^)-Ha antimicrobial peptide may be an interesting and promising option in the substitution of conventional antibiotics.

## INTRODUCTION

*Salmonella* is a bacterium of public health importance due to the high risk of transmission, which can contaminate food or spaces, mainly by its common host, the poultry (Renu et al., 2020). The frequent transmission, as well as the attempt to control *Salmonella*, allowed this microorganism to generate or acquire resistance to several commercial drugs. *Salmonella* accelerated proliferation would be related to the immunity alteration by stress factors, product of excessive manipulation or environmental conditions (Gaggìa et al., 2010). In the last update of the WHO, in 2017, it was declared and published a list of global priorities of resistant bacteria to antibiotics, in which *Salmonella* sp. fluoroquinolone-resistant was classified in the high priority group (number 2). Therefore, based on public health policies, there is a very high degree of concern about these aggressive pathogenic bacteria species. In this scenario, antibiotics used in the poultry industry are increasingly restricted and discovery of new drugs is becoming essential and urgently demanded (WHO, 2017; Tacconelli et al., 2018).

In recent years, antimicrobial peptides (AMPs) have been a central object of study, attributable to their great capacity to control bacterial pathogens, including viruses and fungi (Ahmed et al., 2019; Lin et al., 2019). Specifically, the Ctx(Ile^21^)-Ha antimicrobial peptide is an amphipathic and cationic peptide, isolated from an Brazilian amphibian skin (*Hypsiboas albopunctatus*) (Ferreira Cespedes et al., 2012), which has high antimicrobial capacity, demonstrated in pathogens of public health interest (Vicente et al., 2013). These AMPs are considered natural antibiotics, as they are part of a biological defense innate immune system. Also, AMPs are biocompatible, can modulate immune systems and have high biological activities with minimal concentrations (Ghibaudo et al., 2016). An interesting feature of AMPs is that they can generate a minimal level of bacterial resistance (Malekkhaiat Häffner and Malmsten, 2019). As a result of these attractive characteristics, AMPs application in poultry as a feed additive is promising (Kogut et al., 2013; Roque Borda et al., 2020, 2021b; Silveira et al., 2021)

However, use of these molecules is limited due to instability factors, such as denaturation or acid hydrolysis degradation, produced by gastric acids in the stomach of monogastric animals. To overcome these issues, coated bioformulations to protect bioactive molecules are demanded. Microencapsulation, a standard pharmacotechnical methodology and very established technique in literature, is used to control, protect and maintain compounds’ biological activities (Plazola-Jacinto et al., 2019).

Some types of encapsulations were developed to improve poultry production. For example, spray drying is employed to microencapsulated probiotics and maintain 90% of stability, allowing them to be installed in the chicken intestine (Harimurti et al., 2019). Enteric coating is a protection method widely used in the pharmaceutical area, which permits the targeted transport of biomolecules or drugs to be released at a specific site, depending on the conditions of the polymer used, such as hydroxypropyl methylcellulose phthalate (HPMCP) (Son et al., 2016). This is a modified polymer, derived from cellulose and pH dependent, which it tends to dissolve in liquid solutions at pH > 6.5, playing an excellent role as a drug carrier against intestinal pathogens (Momoh et al., 2020).

These parameters allowed us to design an innovative product based on microencapsulates and enteric coating of a biocompatible molecule with potential antimicrobial activity. The Ctx(Ile^21^)-Ha AMP was chosen due to its properties previously reported by our research group (Vicente et al., 2013). Moreover, a patent described the present developed methodology was deposited at the National Institute of Intellectual Property (INPI BR1020200220489), which is protected throughout the Brazilian territory (Vicente et al., 2020). In this way, the objective of this study was to evaluate the *in vivo* effect of the HPMCP-coated microcapsules containing the Ctx(Ile^21^)-Ha antimicrobial peptide application against *Salmonella* Enteritidis in chickens, to demonstrate its great potential as an innovative natural feed additive in poultry production.

## MATERIAL AND METHODS

### Chemical reagents

HPMCP (Grade HP-55, Nominal Phthalyl Content 31%) was kindly donated by Shin-Etsu Chemical (Tokyo, Japan), and the other chemical reagents were obtained in HPLC grade (Sigma-Aldrich Co., St Louis, MO, USA). *N,N*‐dimethylformamide (DMF) was purchased from Neon Comercial (São Paulo, Brazil), dichloromethane (DCM) was purchased from Anidrol Products Laboratories (São Paulo, Brazil), sodium alginate with low molecular weight (12,000 – 40,000 g mol^−1^, M/G ratio of 0.8) and aluminum chloride were obtained from Êxodo Científica (São Paulo, Brazil). Fmoc-amino acids were purchased from AAPPTEC (Louisville, KY, USA) and other reagents were obtained in HPLC/analytical grade from Sigma-Aldrich (St Louis, USA). Brain Heart Infusion (BHI) broth, Mueller Hinton (MH) agar, Bright Green Agar (BG), selenite broth (SB), nutrient broth (NB), and others microbiological reagents were purchased from SPLABOR (São Paulo, Brazil).

### Ctx(Ile^21^)-Ha antimicrobial peptide synthesis

The antimicrobial peptide Ctx(Ile^21^)-Ha was synthesized manually using solid phase peptide synthesis (SPPS) with Fmoc strategy protocol. The complete methodology is described according to Roque Borda et al. (2021a). Briefly, peptide was assembled at 0.2 mmol scale on a Fmoc‐Rink Amide resin of 0.68 mmol g^-1^ substitution, using three-fold excess and preconditioned for 15 min in DMF and DCM as main SPPS solvents. 4-methylpiperidine/DMF (1:4, v/v) was used to remove the Fmoc amino group protectors from amino acids. Finished the entire peptide primary sequence, Ctx(Ile^21^)-Ha peptide was separated from the resin using a solution containing trifluoroacetic acid/ultrapure water/triisopropylsilane (95:2.5:2.5, v/v/v), at 160 rpm for 2 h at room temperature. Then, samples were freeze-dried (Liotop model K108, Brazil) to obtain the peptide in a white and flocculent powder material.

Peptide purity degree was determined by analytical HPLC (Shimadzu, model Prominence with membrane degasser DGU-20A5R, UV detector SPD-20A, column oven CTO-20A, automatic sampler SIL-10AF, fraction collector FRC-10A and LC-20AT dual-pump, C18 column) at flow rate of 1 mL min^-1^ and detection at wavelength of 220 nm, using as mobile phases 0.045% aqueous TFA (eluent A) and 0.036% TFA in acetonitrile (eluent B) for 30 min. Subsequently, samples were lyophilized and stored until the usage. The Ctx(Ile^21^)-Ha peptide was employed only if the purity degree was higher than 95%. After that, the peptide was confirmed and characterized by ESI-MS (Electron Spray Injection Mass Spectrometry), employing a mass spectrometer (Amazon, Bruker Daltonics, Billerica). Pure Ctx(Ile^21^)-Ha peptide concentrations were determined by UV spectroscopy, considering tryptophan extinction coefficient of 5.600 M^-1^ cm^-1^ at a wavelength of 220 nm.

### *In vivo* analysis

#### Development of Ctx(Ile^21^)-Ha coated-microcapsules (ERCtx)

Ctx(Ile^21^)-Ha was encapsulated by ionotropic gelation method, following the method described in Roque Borda et al. (2021a). Summarily, the peptide-alginate solution was prepared with an initial concentration of 32 (PEP1) and 64 mmol L^-1^ (PEP2) of Ctx(Ile^21^)-Ha peptide in 2% (w/w) sodium alginate homogenized using an UltraTurrax-T18 (IKA-Labortechnik, Germany) at 25,000 rpm min^-1^ and sonicated with an ultrasound probe (Hilscher, Germany) for 15 min. Therefore, a crosslinking solution was prepared with 5% aluminum chloride. Capsules were obtained using a syringe pump (NE-1000, New Era Pump System Inc., USA) with feed flow rate of 1.5 µL h^-1^ at room temperature. After that, they were dried and stored in darkness.

Ctx(Ile^21^)-Ha microcapsules were coated by fluidized-bed method, preparing a coating solution with 10% w/w HPMCP, 25% w/w ammonium hydroxide, 2.5% w/v triethylcitrate and 62.5% of water. The microcapsules were placed on a fluidized-bed (LabMaq MLF 100, Brazil) at 40 °C, 0.25 L min^-1^ blower, 0.4 mL min^-1^ peristaltic pump and 100% vibration as a system condition (Figure S2). The final products (ERCtx) used for *in vivo* evaluation are represented by PEP1 and PEP2. All the microencapsulation development and characterization are described according to Roque Borda et al. (2021a).

#### *In vivo* experiment in chicks

Animal experiments were approved by the local Animal Ethics Committee School of Sciences and Engineering, UNESP, Tupã, Brazil (Proc. n. 06/2018 CEUA). The mortality rate of the chicks was the guiding variable for the calculation of the sample size. Although, due to the absence of similar studies for parameters estimation necessary for the calculations, the minimum number of chicks required to demonstrate the efficacy of the treatment in reducing mortality was not determined. However, the degree of invasiveness present in this study, together with the ethical requirements in the use of animals in experiments, added to the preservation of the quality of handling, led to the use of 45 animals per treatment, totalizing 135 animals.

This number was also in agreement with Montgomery and Runger (2012), when they report on the minimum conditions to use the approximation of the binomial by normal distribution, which is necessary when calculating the sample size that involves proportional estimation. The approximation conditions are: *np* > 5 and *np*(1-*p*) > 5, where “*n*” is the sample size, “*p*” is the proportion of the event of interest: in this case, the death of the chicks. These same authors note that, in general, the better approximation is given for large samples (*n* > 40).

Therefore, to perform the *in vivo* assays, 135 commercials chicks from Hy-lines Brown, Brazil, were acquired from a commercial hatchery. Chick swabs were taken at random, and a box swab sample, to detect *Salmonella* Enteritidis (*S*. Enteritidis) in newborn chickens and verify that they were free of infection. Thus, confirming that all the chicks used in this experiment were negative for this bacterium, the samples were cultured in SB 2X for 24 h at 37 °C.

For the inoculum, *Salmonella* Enteritidis resistant to nalidixic acid and spectinomycin (SE Nal^R^Spc^R^, code P125109), bacterial strain from donated by the Laboratory of Ornithopathology FCAV/UNESP, was used. *S*. Enteritidis was grown in NB for 24 h at 37 °C. All chicks challenge was carried out with using 0.2 mL of 10^9^ CFU mL^-1^ of *S*. Enteritidis.

Chicks were randomly distributed into three groups, separated in 45 chicks for each treatment. They were identified with enumerated tape around the right leg. Since the first day of the experiment, animals received water and feed provided *ad libitum* and doses of antimicrobial peptide Ctx(Ile^21^)-Ha microencapsulated were added to the feed and administered to chicks from the first day of life. Control treatment (CTRL) was defined as that which received only the initial commercial feed for chicks without any additives; the PEP1 treatment received the ERCtx (3.2 µg of Ctx(Ile^21^)-Ha per kg of poultry feed) and the PEP2 treatment received the ERCtx (6.4 µg of Ctx(Ile^21^)-Ha per kg of poultry feed), both microparticles added to the initial commercial feed of the control treatment (isoproteic and isoenergetic for chicks in the first 28 days of life).

For the chick cloacal swab, 15 chicks were selected for each group. A collection of the fecal excretion was performed two times each week. The collected samples were incubated in 3 mL of SB and Novobiocin (Nov) at 37°C for 24 h, to later be seeded in BG Nal/Spec and incubated again. This procedure was repeated throughout the experiment. The results were expressed as presence/absence of *S*. Enteritidis, depending on being positive or negative for the pathogen, respectively (Berchieri A. et al., 2001; Freitas Neto et al., 2007). In addition, chicks were weighed alive from 12 days of age until the end of the experiment.

For the evaluation of intestinal infection, five chickens from each group were sacrificed for the count of *S*. Enteritidis in the cecal content, carried out on days 2, 5, 7, 14, 21 and 28 postinfection (dpi). The samples were collected aseptically, with the help of sterilized forceps and individual scissors for each chick. The previously weighed tubes were conditioned in PBS pH 7.4 in the ratio 1:10, w/v, and were homogenized in vortex. The samples were seeded and cultured on a BG Nal/Spec agar plate at 37°C for 24 h and counted in colony forming units (CFU).

### Statistical analysis

To perform the total count of *S*. Enteritidis in CFU/mL, the data was transformed into a Napierian logarithm (Ln) to adapt the model recommended in ANOVA. Mortality was analyzed using the Chi-square test. The results were analyzed by software R.

## RESULTS

### Peptide analysis

In the initial analysis, 590 mg of crude mass of peptide was obtained which was subsequently purified. The purification yield was 20% with a total mass of 120 mg. The analysis was carried out by HPLC and Mass Spectrometry, confirmed the obtaining of Ctx(Il e^21^)-Ha antimicrobial peptide (MW = 2,289.72 g mol^-1^) shown in the Supplementary Material (Figure S1).

### *In vivo* results

#### Post-inoculation mortality

The mortality results showed significant (α = 0.05) influence of the application of the Ctx(Ile^21^)-Ha microparticles in the treatments on the registered mortality percentages (P=0.03), by using Qui-square test, with 2 degree of freedom (df = 2). Therefore, mortality percentages differ significantly between treatments. Table 1 allows the estimating the risk of death of the chicks according to the treatment, which are, in fact, conditional probabilities. There are three risks that can be estimated from Table 1.

In control treatment (CTRL), the estimated risk of death for a chicken (R_CTRL_) corresponds to the probability estimation of chicks death, given the non-ingestion of the microparticles with antimicrobial peptide Ctx(Ile^21^)-Ha, that is, 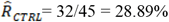 In parallel, risk of death for a hen treated with PEP1, which is the estimate of the probability of death of the chick given the ingestion of (32 mmol L^-1^ peptide is 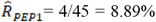 Finally, the risk of death for a hen treated with PEP2, which corresponds to the estimated probability of death of the chick given the ingestion of (64 mmol L^-1^ peptide, that is, 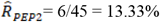

The mortality results were explored with the percentage distribution conditioned by each treatment, where the proportion of the results was presented as a function of the corresponding treatment to which the chicks were subjected (Figure 1). Due to the statistics illustrated in Figure 1, risks of death were compared two by two, using estimated Relative Risk 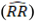, a statistic that quantifies the relationship of higher mortality through the relationship between risks (González et al., 2019), being the numerator the highest risk and the denominator the lowest risk.

**Figure 1.**
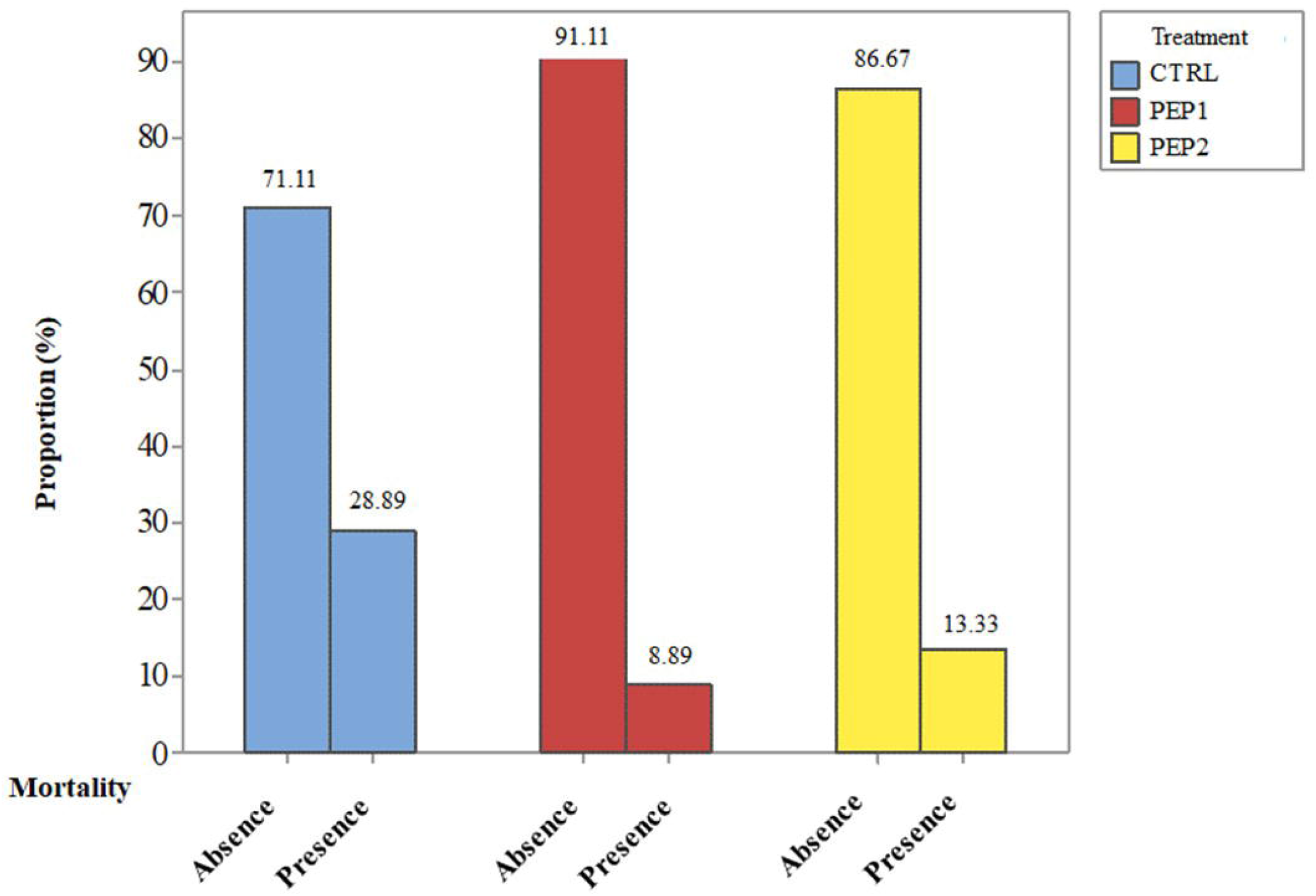
Percentage distribution of mortality due to the *in vivo* treatments analyzed.

In this study, PEP1 and PEP2 treatments have the function of protecting the chicks from the direct and indirect harmful effects of *S*. Enteritidis inoculation. Therefore, control group corresponds to the exposure group and, consequently, of higher risk. Thus, three 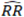 estimates could be produced, but only two were of real interest for analysis.

The first was the RR of mortality among the animals in the CTRL and PEP1 treatments 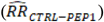:

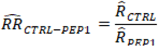

According to the results, a value of 3.25 (P = 0.01) was obtained. This implies that the risk of death of the hen is 3.25 times higher in the CTRL condition compared to PEP1.

The second, the estimated relative mortality risk between animals in the CTRL and PEP2 treatment 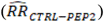:

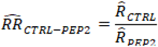

and the following results were obtained, a value of 2.17 (P = 0.04) is reached, which indicates that the risk of death of the hen is 2.17 times higher in the CTRL condition in compared to PEP2.

Importantly, the use of the hypothesis test (H_0_ or H_1_) performed was one-sided since the treatment is unlikely to increase mortality, at 5% of significance level (α = 0.05). Thus, the null hypothesis (H_0_: RR = 1) is rejected, in both risk tests (P = 0.04) in favor of the alternative hypothesis (H_1_: RR > 1), which allows to use the 95% Confidence Interval (CI) of Relative Risks as a form of interval estimation for the RR population. This is represented by:

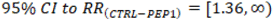

That is, with 95% CI, it is possible to affirm that the true RR in question (the population) is at least greater than or equal to 1.36. This implies that the true mortality in a population is at least 36% higher for control group animals compared to PEP1 treated group:

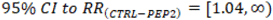

That is, with 95% CI, it is possible to affirm that the true RR in question (the population) is at least 1.04. This implies that the true mortality in a population is at least 4% higher for the chicks in the control group, compared to the PEP2 treated group.

In this experiment there was no reduction in mortality when the peptide concentration was increased. Therefore, it is not necessary to test the statistical difference in mortality risk between these two doses; Also, a higher protection (lower mortality) was obtained with fewer resources (peptide mass), which is important to an industrial approach. That is, due to the results, the reduction in mortality does not improve due to the increase in concentration. However, in the best of cases, it remains the same. Furthermore, it is highlighted that there is statistical evidence that PEP1 treatment reduces total mortality, and not only due to *S*. Enteritidis infection. They can be used to establish a metric that quantifies the protection acquired by chicks, because they were also subjected to a treatment with peptides (PEP1 or PEP2). In addition, this can be due to the nature of the action of antimicrobial peptides, which is to protect the chicks against *S*. Enteritidis (results shown in Table 1 and Figure 2).

**Figure 2.**
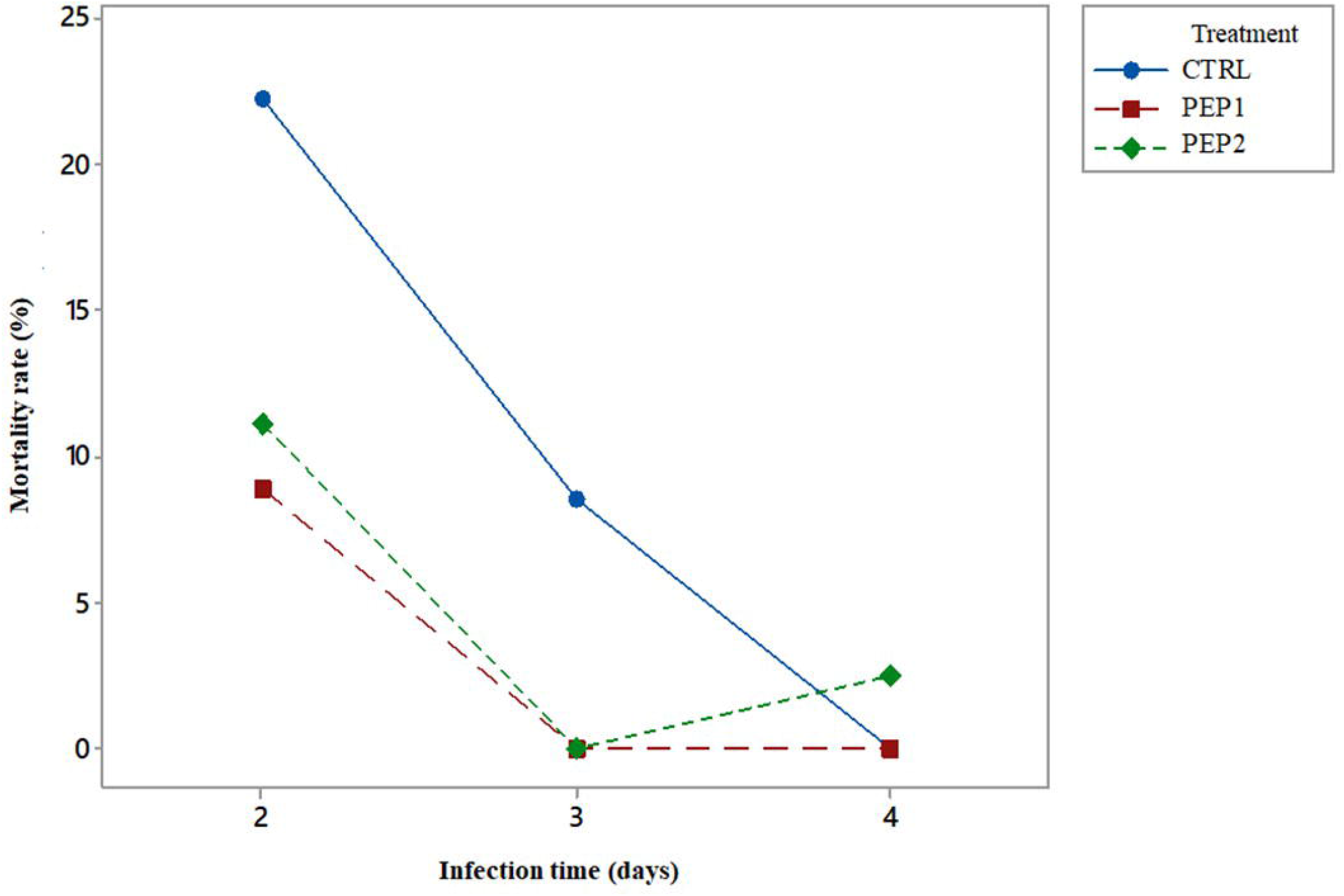
Mortality rate as a function of time of infection.

Protection Factor (PF) is the statistic that quantifies the reduction in mortality risk due to the use of PEP1 or PEP2. In this case, mathematically, the PF is nothing more than the opposite of RR, that is, PF = 1 − (RR^-1^). Therefore, the estimation of the Protection Factor that PEP1 treatment has on mortality, when compared with the CTRL is given by:

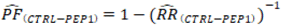

Therefore, 1 − (1/3.25) = 0.69. Thus, it is specifically estimated that treatment with PEP1 reduces the risk of death of chicks by 69% (P<0.01), compared to the CTRL group. When using the one-sided interval estimation, with 95% confidence, for 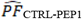:

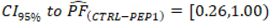

This implies that the reduction in mortality from the use of PEP1 is at least 26%, compared to the control group. The estimate of the protection factor exerted by PEP2 treatment on the control group, and is given by:

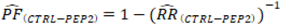

Therefore, 1 − (1/2.17) = 0.54. This result indicates that PEP2 treatment reduces the risk of mortality by 54% compared to the CTRL group (P = 0.04). Using the unilateral interval estimate for 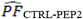 with 95% confidence:

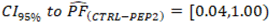

Consequently, the reduction in mortality from the use of PEP2 is at least 4% (more precisely, 4.05%), compared to the CTRL group.

The mortality rate after 5 dpi (days post-infection, 7th day of life) was zero for the chicks, subjected to all the treatments studied. Thus, critical analysis period was concentrated from 2 dpi to 4 dpi day of the experiment. With clear evidence, chicks treated with the ERCtx had a mortality rate of zero at 3 dpi (Figure 2). In relation to PEP2, there was only one death at 4 dpi, a fact that differs from the trend shown by treatments, and that may be caused by some eventuality, but it is not possible to conclude with certainty. There is a significant difference between the mortality rate of the CTRL group compared to the PEP1 and PEP2 treatments, on the second and third days of infection (P <0.05). This shows that the antimicrobial peptide has the effect of reducing the risk of mortality already at the beginning of infection, when chickens ingest the microencapsulated peptide, which corresponds to the most acute phase of their mortality.

#### Using Power Test to analyze results

The one-sided 95% CI showed that the reduction, in percentage points, in the mortality rate from the control group to the PEP1 treatment group would have the lower limit (LL = 0.07 = 7%). This means that the existing population reduction would be at least 7%, which is highly satisfactory. For the reduction from CTRL to PEP2 it would be with LL = 0.02 = 2%, which is less than satisfactory. For this reason, it is not necessary to increase the sample size to demonstrate the efficacy of the Ctx(Ile^21^)-Ha antimicrobial peptide, when considering the performance of PEP1 treatment in reducing the mortality rate.

When assessing the sufficiency of the sample size, analysis with the Power Test can be included, which corresponds to the sensitivity of the test to reject the null hypothesis unequivocally, which can further improve test performance (Roush and Tozer, 2004). For this reason, a simulation was developed for different mortality rates lower than those obtained in the control treatment, according to the alternative hypothesis. The simulation for the sample size calculations was based on the estimates obtained, with a significance level of 5% and power test of 80%, considered by Cohen (1988) as acceptable. In Figure 3, the minimum number of sample units per treatment (for example, number of chicks used) is shown for the test to detect the reduction in mortality for proportions of 25 to 5%.

**Figure 3.**
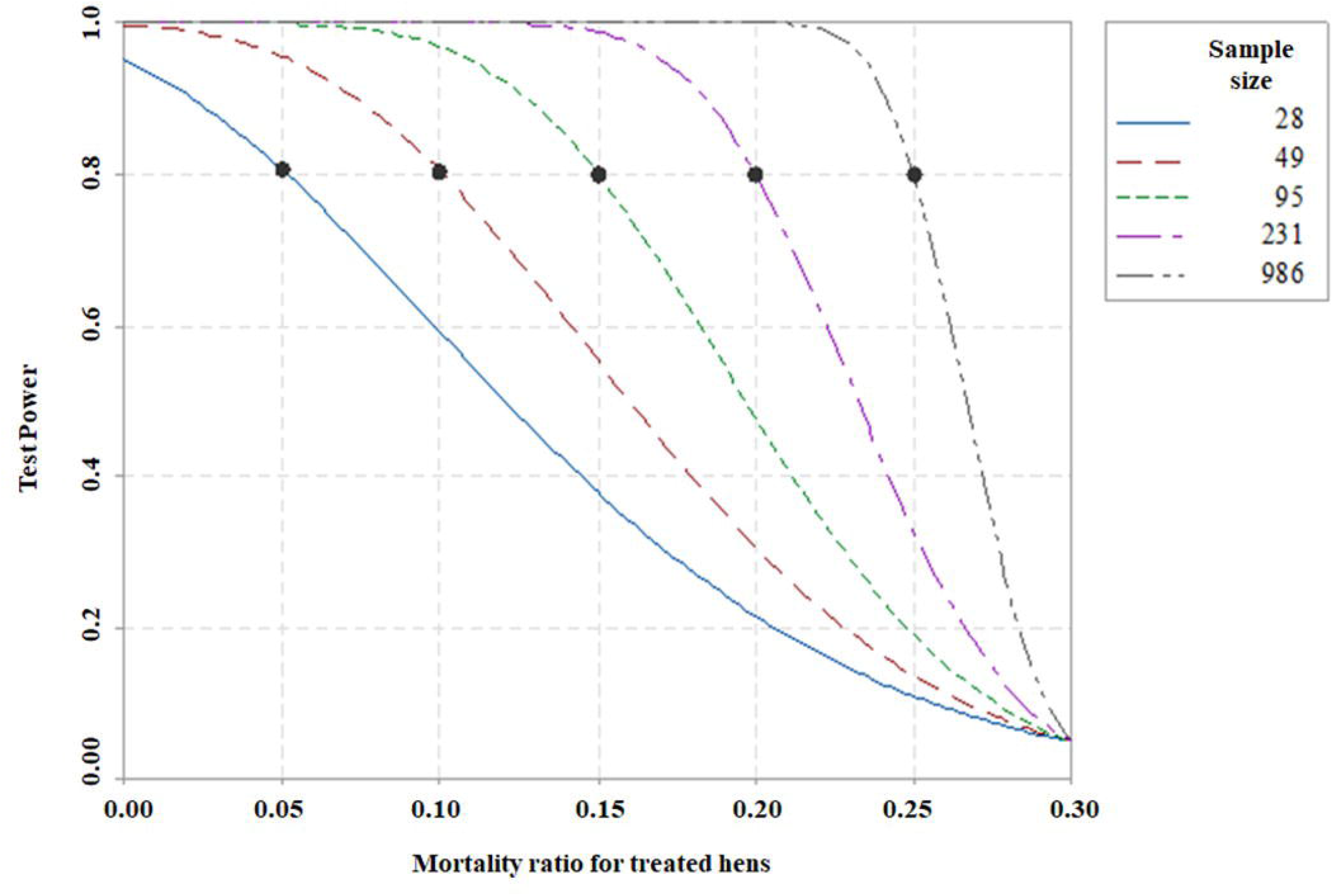
Calculation of the theoretical sample size to demonstrate the test power obtained in this experiment.

According to Figure 3, it is evident that the more the mortality proportion is reduced, the smaller the sample size necessary to detect the decrease in mortality, with a power of 80% and a significance of 5%. Mortality rates between 5 and 10% are known to require between 28 and 49 chicks per treatment, respectively. This result affirms that the 45 chicks used per treatment in this experiment would be sufficient to detect a reduction in mortality in the control treatment of the order of 20 percentage points or more, which occurs in the treatment with the lowest dose of peptide (PEP1). In another situation, if it is considered only for comparison (CTRL for PEP2), then a significance level greater than 5% would be adopted, or a power less than 80%, which could compromise the sensitivity of the test, or in general, test another four animals for each treatment to complete 49 animals per treatment, which would be unnecessary at this time, given the results found.

#### Cecal content

The data for the *S*. Enteritidis count in CFU mL^-1^ required transformation to a natural logarithm (Ln), to adapt the recommended model in ANOVA. The F test for treatment purposes showed statistical evidence (P <0.02) that there is a difference between treatments. Likewise, the time effect of infection on the Ln count was strongly significant (P = 0.00), specially on the second day of infection (5 days of life). In addition to the main effects, the interaction effect was also investigated (P = 0.24).

This result shows that there is no evidence of an interaction between treatment and age, that is, the treatment showing the best performance is still PEP1, especially on the second day of infection, and after that this treatment alone does not differ significantly from the others, as shown in Figure 4.

**Figure 4.**
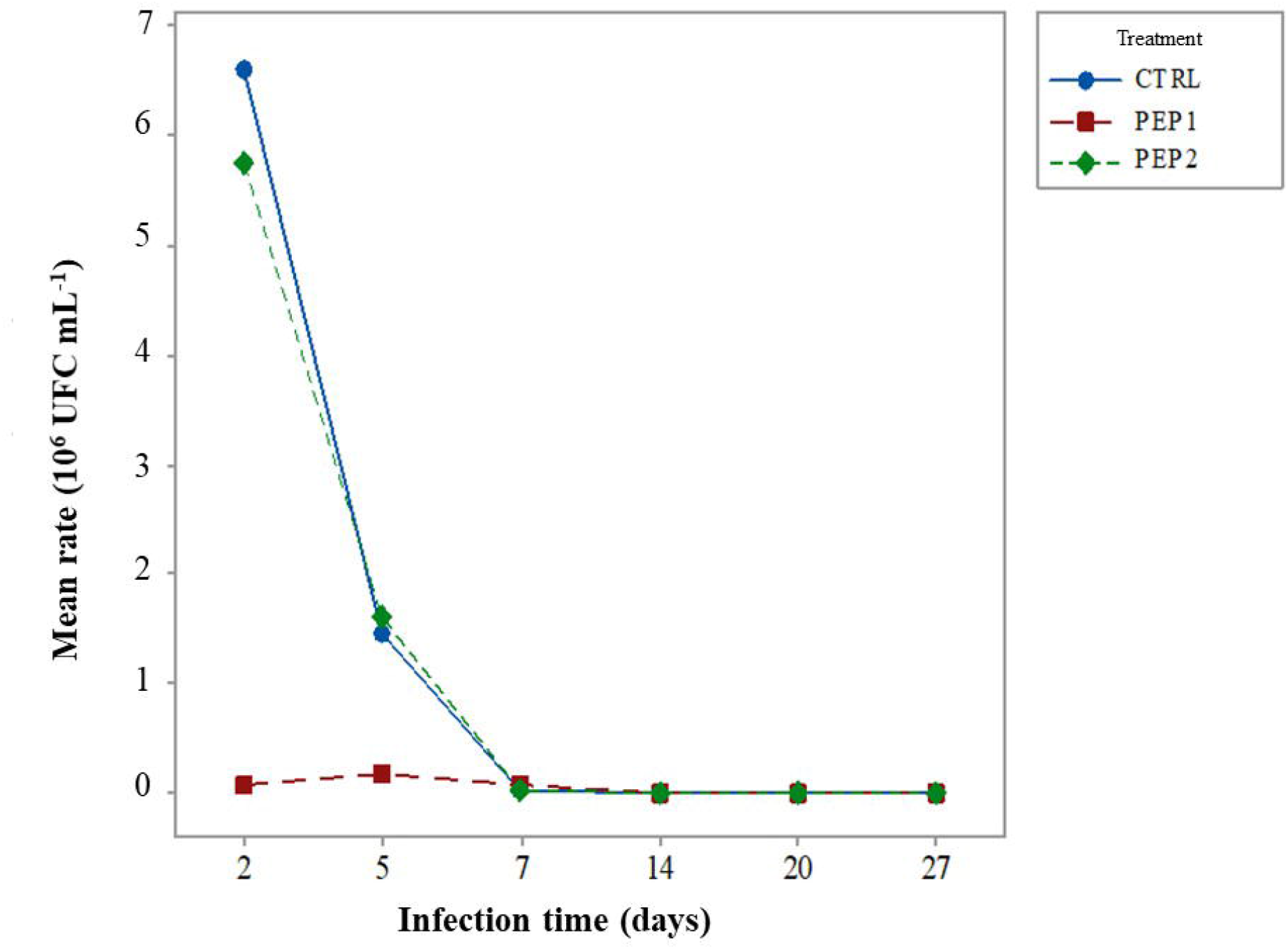
Mean count rate of *S*. Enteritidis, which is dependent on the treatments and time of infection.

After ANOVA, a comparison test of the control treatment (CTRL) was performed against the other treatments (PEP1 or PEP2) using Dunnett’s test, which, in turn, showed statistical significance only between CTRL and PEP1 (P<0.05).

#### Chick cloacal swab

Results of the follow-up of the infection with swab method of the *S*. Enteritidis inoculum (Figure 5) at a concentration of 10^9^ CFU mL^-1^, were subjected to the Chi-square test (P>0.05), where the *p*-value found was 0.88. This result indicates that there is no evidence that the proportion of presence or absence is different between treatments. Therefore, based on this sample, it cannot be said that the treatment influences the absence or presence of the inoculum.

**Figure 5.**
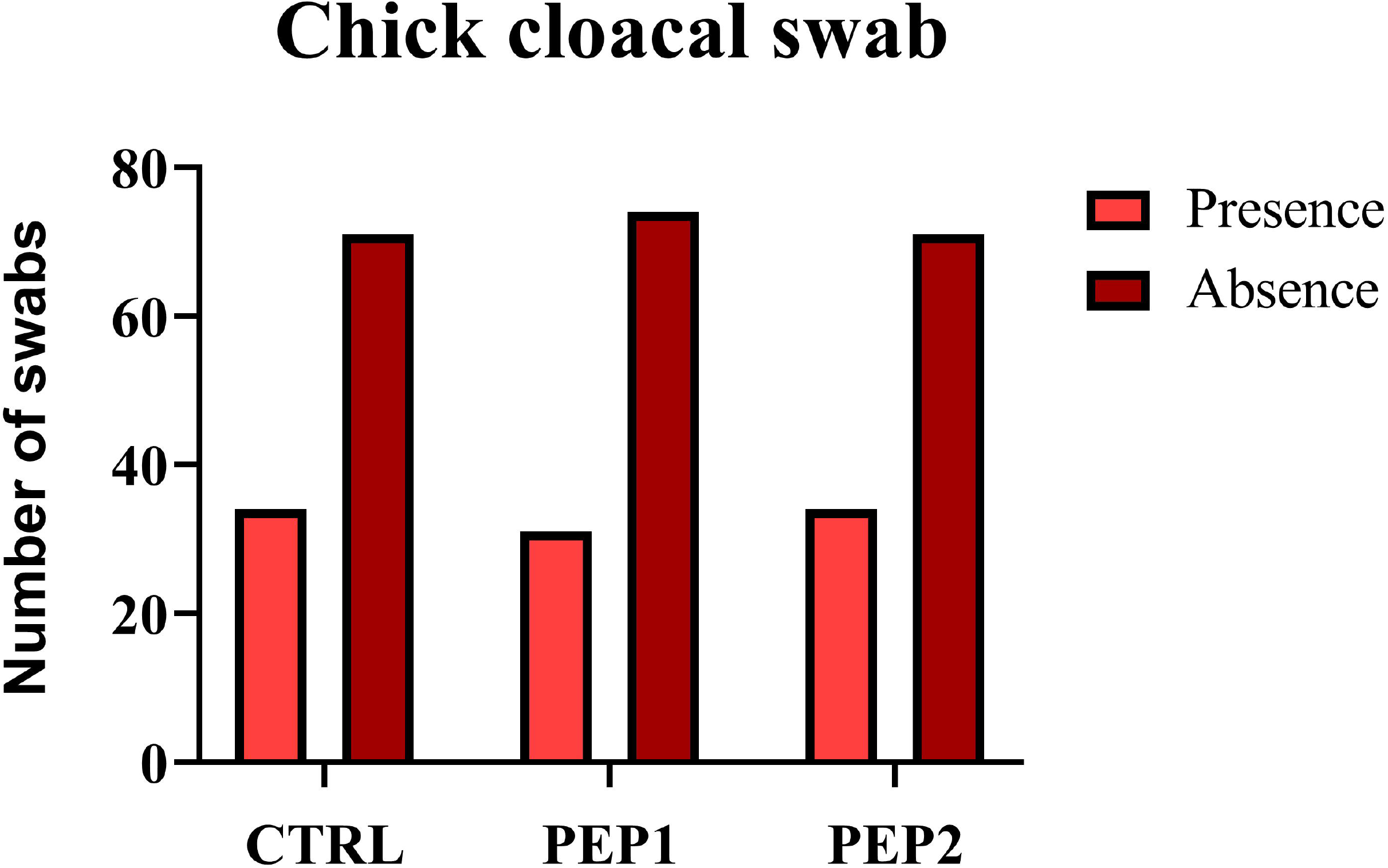
Evaluation of the effect of antimicrobial peptide on *Salmonella* Enteritidis fecal excretion during 28 days.

#### Weighing of the chicks

The mean and standard deviation of the results obtained from weighing during the experiment are shown in Table S1. From the inferential perspective, the F-test for treatment purposes showed moderate evidence (P = 0.07) that there is a difference between treatments. The effect of age on weight, on the other hand, was strongly significant (P = 0.00), due to the intrinsic development of the body mass of the chicks, especially in the first weeks of life. In addition to these main effects, the presence of an interaction effect was also investigated (P = 0.74). This result shows that there is no evidence of interaction between treatment and age, that is, the effect of the treatments does not depend on age. These results could be illustrated using boxplot (Figure 6).

**Figure 6.**
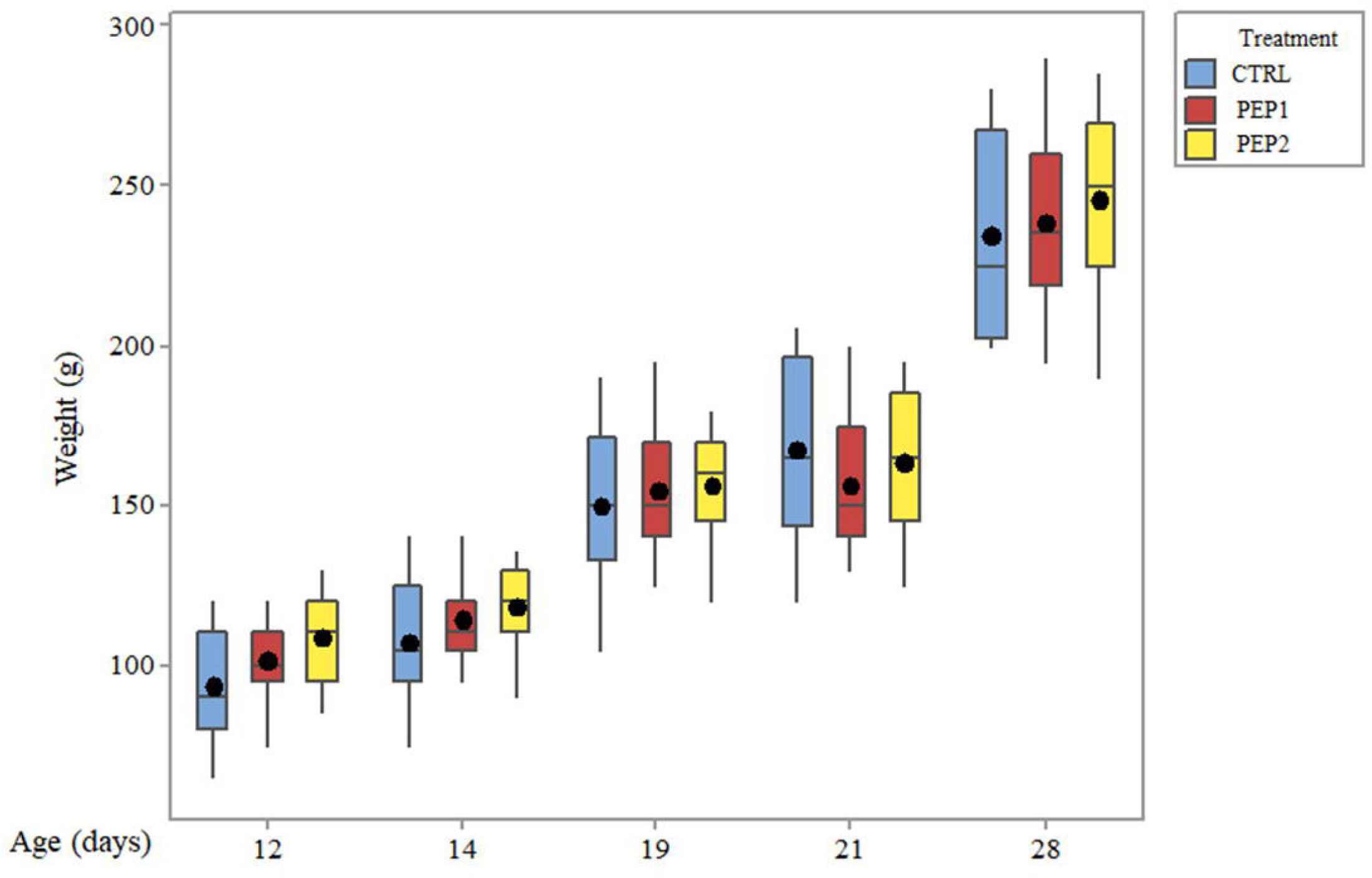
Weight distribution according to the treatments and infection time, time to which the chicks were subjected.

## DISCUSSION

It is known that the mortality rate caused by *S*. Enteritidis is low (Barbosa et al., 2017). However, infection with this bacterium weakens the immune system and causes collateral damage that affects nutrient absorption (He et al., 2005). Consequently, other bacteria can act as opportunists, colonizing the intestine and causing poultry death. Systemic poultry infection would be linked to the influence of the flagella in some *Salmonella* sp. serovars (Barbosa et al., 2017). Thus, when it comes to newborn chickens (up to 5 dpi), some studies suggest the use of immune stimulators to reduce mortality rate (He et al., 2005). Another study affirms that the BT peptide was able to promote mRNA transcription for Toll-like receptors (TLR), responsible for producing a pro-inflammatory response with cytokines and, consequently, activating the immune response. They also highlighted that the use of AMPs in the first 4 days of life is important, and that its best application would be orally (Kogut et al., 2013).

Recent study indicates that AMPs have the same bacteriostatic potential compared to conventional antibiotics, and they also increased the content of white blood cells, making them an excellent replacement alternative for bio-sustainable poultry production, even better than other natural components (Xu et al., 2021). A study with essential oils in a combination of *Syzygium aromaticum* and *Cinnamomum zeylanicum* showed antimicrobial activities against *S*. Enteritidis and *S. typhimurium* (0.322 - 0.644 mg mL^-1^ and 0.644 - 1.289 mg mL^-1^, respectively). However, these MIC values were very low compared (9.32 and 37.30 µg L^-1^, respectively) to those reached by Ctx(Ile^21^)-Ha antimicrobial peptide (Roque Borda et al., 2021a).

MccJ25 is a highly studied cyclic recombinant AMP, presenting a broad bactericidal spectrum, mainly against *Salmonella* sp. (Vincent et al., 2004). This AMP showed that its function is not only to eliminate the bacteria and improve the fecal microbiota, but also influence intestinal morphology by improving texture and reducing inflammation after infection (Wang et al., 2020). Also, bacteriocins are an AMP group studied for use in poultry and swine industry. They are produced by some bacteria (the majority by Gram-positive) and present interesting effects by reducing the content of pathogenic bacteria, such as *Salmonella* sp. and *Campylobacter jejuni* (Ben Lagha et al., 2017). Other bacteriocins were studied to reduce *Salmonella* in broilers using a dose of 2.5 g k^-1^, exhibiting an increase in the weight of the chickens and a slight bacterial decrease (Wang et al., 2011). Unlike our results, which could be due to the minimum dose used in this experiment, which is thousands of times less than the reference, our work used µg and the reference used mg of peptide. However, other research indicates that application of swine intestine (Wang et al., 2009) as a food supplement influences positively broilers’ weight, as well as increased villus height (Liu et al., 2008). For explanation, the hypothesis is if Ctx(Ile^21^)-Ha peptide concentration is incremented, it will be possibly to visualize a more pronounced increase in weight in chicks. Although, for a pilot *in vivo* experiment, the Ctx(Ile^21^)-Ha peptide microcapsules showed very promising and interesting results.

Based-studies of previous AMPs-encapsulates, bacteriocins were protected using polyvinylpyrrolidone as encapsulating material showed promising results, due to decrease content of *C. jejuni*. However, the AMP dose employed was very high compared with our encapsulated products (500 mg of AMP per kg^-1^ of poultry feed) (Stern et al., 2005). Our product is encapsulated based on sodium alginate, which is a cheap and more biocompatible polymer (Lee and Mooney, 2012). Sodium alginate encapsulations loaded with specific phages (f3αSE) for *S*. Enteritidis have been made, achieving up to 80% protection of viability of this product at gastric pH (Soto et al., 2018). This confirms and corroborates that our developed microcapsules can satisfactorily contain and protect the Ctx(Ile^21^)-Ha AMP, as they also have an additional protective layer with HPMCP, which is only degraded at intestinal pH.

Consequently, this study showed a decrease in mortality rate in first days of life, which certifies the success of the encapsulated and coated antimicrobial peptide. Values of anti-*S*. Enteritidis activity *in vivo* did not have a significant difference. However, there is an interesting result in first days of life, decreasing total count. This could be due to the relationship between the amount of microcapsule ingested and the total volume of the intestine, and that each time the chicks grew, this relationship would be more different. Therefore, it is suggested that further experiments employing higher doses can be performed to achieve a total bacterial decrease.

## CONCLUSIONS

*In vitro* studies revealed that the microencapsulated Ctx(lle^21^)-Ha peptide presented antimicrobial activity with pathogens from poultry sector such as *Salmonella* Enteritidis, *Salmonella typhimurium* and *Escherichia coli*. Results of the *in vivo* analyzes allow to conclude that the antimicrobial peptide Ctx(lle^21^)-Ha presented positive, significant and promising results in relation to the reduction of younger chicken’s mortality and the bacterial count. Mainly for PEP1 treatment, where there is a 69% reduction in the risk of death. Regarding the weight of the chickens, in two doses of antimicrobial peptide used, there was a significant difference between treatments and this result shows that there is no evidence of interaction between treatment and age, that is, the effect of the treatments does not depend on age. Finally, it is concluded that there is a potential effect of the microencapsulated-coated antimicrobial peptide Ctx(lle^21^)-Ha in poultry, which enable the application of the peptide using very low mass, compared to other studies in literature.

## Acknowledgments

This work was developed with the support of São Paulo Research Foundation/FAPESP (Process number 2016/00446-7), master (Process number 2018/25707-3) and undergraduate scholarships (Process number 2017/21822-0). We thank the technical assistants of Laboratory of Chemistry and Biochemistry from São Paulo State University (Unesp), School of Sciences and Engineering, Tupã; University of Araraquara which made available the use of the fluidized-bed equipment and Shin-Etsu company for gently donate and provide the HPMCP coating for the experiments. Finally, the research group “Peptides: Synthesis, Optimization and Applied Studies PeSEAp”.

## TABLES

**Table 1.** Distribution of chick mortality by treatment after 4 days of infection.

**Table S1.** Descriptive statistics of weight (g) according to treatments and age.

## FIGURE CAPTIONS

**Figure S1. A**. Chromatographic profile of crude peptide by HPLC at 220nm. **B**. Chromatographic profile of purified peptide by HPLC at 220nm. **C**. Chromatographic profile of crude peptide by LC/MS at 220nm for confirmation.

**Figure S2.** Microcapsules obtained after ionic gelation and fluidized bed.

