## SUPPLEMENTARY MATERIAL for "HPMCP-coated microcapsules containing the Ctx(Ile^21^)-Ha antimicrobial peptide reduces the mortality rate caused by resistant *Salmonella* Enteritidis in poultry"

**Name(s) of Author(s)**

Cesar Augusto Roque Borda^1^, Larissa Pires Pereira^2^, Elisabete Aparecida Lopes Guastalli^3^, Nilce Maria Soares^3^, Priscilla Ayleen Bustos Mac-Lean^2^, Douglas D'Alessandro Salgado^2^, Andréia Bagliotti Meneguin^4^, Marlus Chorilli^4^, Eduardo Festozo Vicente^2*^

**Author Affiliation(s)**

^1^ São Paulo State University (Unesp), School of Agricultural and Veterinarian Sciences, Jaboticabal, São Paulo – Brazil. 14884-900.

^2^ São Paulo State University (Unesp), School of Sciences and Engineering, Tupã, São Paulo – Brazil. 17602-496.

^3^ Poultry Health Specialized Laboratory, Biological Institute, Bastos, São Paulo – Brazil. 17690000.

^4^ São Paulo State University (Unesp), School of Pharmaceutical Sciences, Araraquara, São Paulo – Brazil. 14801-902.

***Address for correspondence:**

*Eduardo Festozo Vicente

São Paulo State University (UNESP), School of Sciences and Engineering, Tupã, São Paulo – Brazil.

**SUPPLEMENTARY MATERIAL**

**RESULTS**

**Peptide analysis**

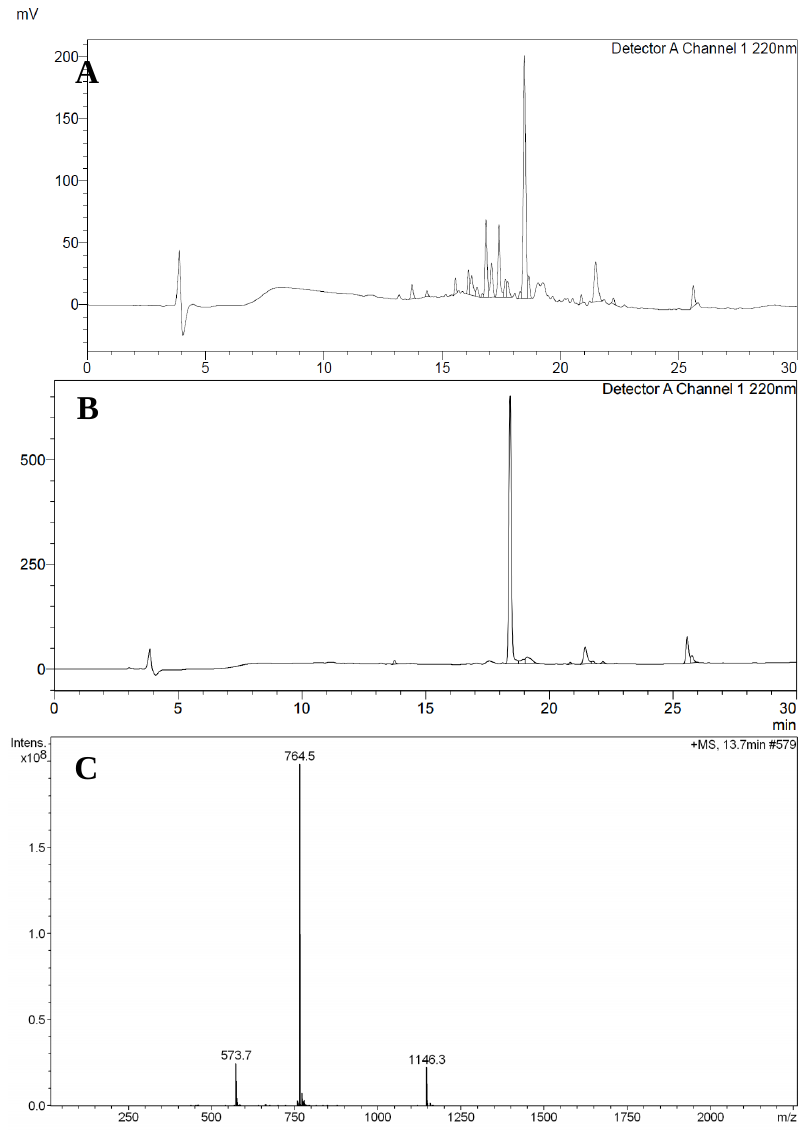

**Figure S1. A.** Chromatographic profile of crude peptide by HPLC at 220nm. **B.** Chromatographic profile of purified peptide by HPLC at 220nm. **C.** Chromatographic profile of crude peptide by LC/MS at 220nm for confirmation.

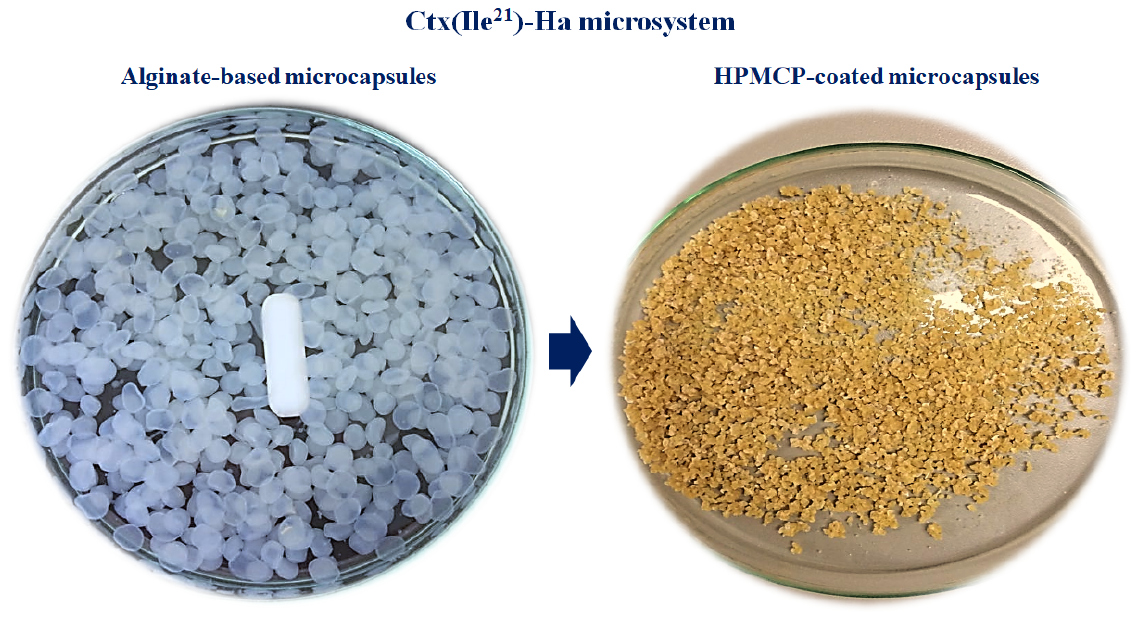

**Figure S2.** Microcapsules obtained after ionic gelation and fluidized bed**.**

***In vivo* results**

**Weighing of the chicks**

**Table S1.** Descriptive statistics of weight (g) according to treatments and age.

| **Age (days)** | **Treatment** | **N** | **Mean (g)** | **SD** | **CV%** | **Min.** | **Q1** | **Median** | **Q3** | **Max.** |
| --- | --- | --- | --- | --- | --- | --- | --- | --- | --- | --- |
| 12 | CTRL | 15 | 93.33 | 16.87 | 18.07 | 65.00 | 80.00 | 90.00 | 110.00 | 120.00 |
|  | PEP1 | 15 | 101.67 | 12.05 | 11.85 | 75.00 | 95.00 | 100.00 | 110.00 | 120.00 |
|  | PEP2 | 15 | 108.67 | 14.20 | 13.07 | 85.00 | 95.00 | 110.00 | 120.00 | 130.00 |
| 14 | CTRL | 15 | 107.00 | 18.40 | 17.20 | 75.00 | 95.00 | 105.00 | 125.00 | 140.00 |
|  | PEP1 | 15 | 114.33 | 11.63 | 10.17 | 95.00 | 105.00 | 110.00 | 120.00 | 140.00 |
|  | PEP2 | 15 | 118.33 | 14.35 | 12.13 | 90.00 | 110.00 | 120.00 | 130.00 | 135.00 |
| 19 | CTRL | 15 | 149.64 | 24.45 | 16.34 | 105.00 | 132.50 | 150.00 | 171.25 | 190.00 |
|  | PEP1 | 15 | 154.33 | 19.72 | 12.78 | 125.00 | 140.00 | 150.00 | 170.00 | 195.00 |
|  | PEP2 | 15 | 156.33 | 18.56 | 11.87 | 120.00 | 145.00 | 160.00 | 170.00 | 180.00 |
| 21 | CTRL | 15 | 167.14 | 26.73 | 15.99 | 120.00 | 143.75 | 165.00 | 196.25 | 205.00 |
|  | PEP1 | 15 | 156.33 | 20.48 | 13.10 | 130.00 | 140.00 | 150.00 | 175.00 | 200.00 |
|  | PEP2 | 15 | 163.33 | 22.81 | 13.96 | 125.00 | 145.00 | 165.00 | 185.00 | 195.00 |
| 28 | CTRL | 15 | 234.40 | 32.30 | 13.76 | 200.00 | 202.50 | 225.00 | 267.50 | 280.00 |
|  | PEP1 | 15 | 238.60 | 28.68 | 12.02 | 195.00 | 219.00 | 235.00 | 260.00 | 290.00 |
|  | PEP2 | 15 | 245.38 | 28.17 | 11.48 | 190.00 | 225.00 | 250.00 | 270.00 | 285.00 |

***Note:*** *N = animal number, SD = standard deviation, CV = coefficient of variation, Q = quartile.*
